# Maternal age alters recombination rate in *Drosophila pseudoobscura*

**DOI:** 10.1101/2020.07.20.212548

**Authors:** Keeley A. Pownall, Hannah N. Taylor, Ulku H. Altindag, Laurie S. Stevison

## Abstract

As individuals senesce, factors correlated with fitness, such as fecundity and longevity, decline. Increased age also alters recombination rates in a variety of taxa. Changes in individual recombination rate, or ‘recombination rate plasticity’, can increase meiotic errors. In *Drosophila melanogaster*, multiple studies on maternal age and recombination rate have found a characteristic pattern where rates initially increase, then decrease, then increase again relative to controls. Here, this phenomenon was investigated in *D. pseudoobscura*. First, fecundity and survivorship were investigated to guide the choice of treatment age. Then, a large-scale recombination analysis (N=23,559) was set up using three X-linked phenotypic markers. Recombination rate differences in two genomic intervals were measured in females aged to 7 days (control) and 35 days (selected treatment age) prior to mating, with progeny collection continuing for 12 days post-mating in 72 hour timepoints. Results revealed a 3.39% increase in recombination rate due to maternal age (p=0.025), for the first 72 hour time point in one of the two marker intervals. For both genomic intervals, recombination rates were higher in the age treatment for the first time point and lower in later time points of the experiment. Next, these data were used to investigate crossover interference, which decreased with maternal age in the first time point and increased in the last time point. Overall, these results suggest that the mechanisms responsible for recombination rate plasticity may differ between maternal age and stressors, such as temperature.

## Introduction

As organisms age, molecular and physiological changes occur that lead to reduced survival and fecundity (Flatt and Schmidt 2009). Parental age has also been shown to be associated with developmental abnormalities in offspring (Parsons 1962, 1964). Due to these apparent negative effects on overall fitness, the existence of senescence has long presented a conundrum to evolutionary biologists (Charlesworth 2000). Because senescence declines more rapidly after reproduction, evolutionary biologists have reconciled the ubiquitous pattern by positing that natural selection is not strong enough to counter the deleterious effects post-reproduction (Charlesworth 2000; Flatt and Schmidt 2009).

One consequence of aging is a subsequent change in meiotic recombination rates. Because crossing over, or synapsis, stabilizes chromosomes during meiosis, rate changes can lead to non-disjunction, and ultimately aneuploidy (Hassold and Hunt 2001). Additionally, recombination generates novel genetic variation, thus changes in rates of crossing over can impact how natural selection either generates or breaks down haplotypes with combinations of beneficial and harmful alleles (Felsenstein 1974; Smith and Haigh 1974).

In 1915, Bridges noticed a difference in rates of crossing over due to age in *Drosophilamelanogaster* (Bridges 1915). Bridge’s work on alterations in recombination rates was among the first to demonstrate what would later become known as ‘*recombinationrateplasticity’* (Agrawal *et al*. 2005; Stevison *et al*. 2017). This phenomenon has since been demonstrated in a variety of organisms and under a multitude of conditions (reviewed in Stevison *et al*. 2017). Although recombination rates have been investigated for influence by both maternal and paternal age in mammals (Kong *et al*. 2004; Vrooman *et al*. 2014), recombination does not occur in male fruit flies (Morgan 1910). Therefore, work in *Drosophila* focuses entirely on females.

In his experiments, Bridges allowed females to lay eggs continuously for 10-days at a time, transferring them to fresh food between time points to create ‘broods’, or clutches, of fruit flies (Bridges 1915). Bridges found a 20% decline in recombination rate as a result of age by comparing the first and second 10-day broods. He further showed that coincidence, a measure of the proximity of crossovers relative to one another, also varied as a result of female age. Similarly, a more recent study in humans concluded that recombination rates increased with maternal age because of the relaxation of crossover interference, resulting in an increased coincidence (Campbell *et al*. 2015).

Later, experiments of multiple regions across the third chromosome revealed variation in the direction of change in recombination rate as a result of age (Bridges 1927). In fact, recombination rates first increased, then declined, followed by a second peak rate, resulting in an overall M-shaped curve (Bridges 1927). This “rise-fall-rise” pattern was observed in later studies of the same chromosomal regions (Redfield 1966), suggesting that the dynamics of maternal age and recombination rate vary spatially along the chromosome, and temporally within an individual. In humans, a positive relationship between recombination rate and maternal age has been observed (Kong *et al*. 2004). However, research on age and recombination rate in *Caenorhabditiselegans* and *Musmusculus* noted a decline in recombination rate with age (Henderson and Edwards 1968; Rose and Baillie 1979). Together, these studies suggest that the direction of how maternal age impacts recombination rates varies among taxa and should therefore be studied more broadly to further our understanding of this complex phenomenon (Parsons 1988; Stevison *et al*. 2017).

Here, we investigate the effect of maternal age on recombination rate in *D. pseudoobscura*. This species of *Drosophila*, which is ∼30 million years diverged from the classic model, *D. melanogaster* (Throckmorton 1975), was the second *Drosophila* species to have its genome completely sequenced and is commonly used for chromosomal studies, which makes it a good model for recombination studies (Hales *et al*. 2015). *D. pseudoobscura* has also recently been investigated as the second *Drosophila* species to show evidence of recombination rate plasticity (Stevison *et al*. 2017). However, an earlier investigation in this species revealed no significant differences due to age on recombination rate (Manzano-Winkler *et al*. 2013). In that study, they used an experimental design similar to the earlier comparisons between clutches (Bridges 1915). The first clutch was allowed to lay eggs for 9 days, with females ranging in age from 1-9 days of age, and the second was allowed to lay eggs for 7 days, with females ranging in age from 10-16 days of age (Figure 1A).

**Figure 1.**
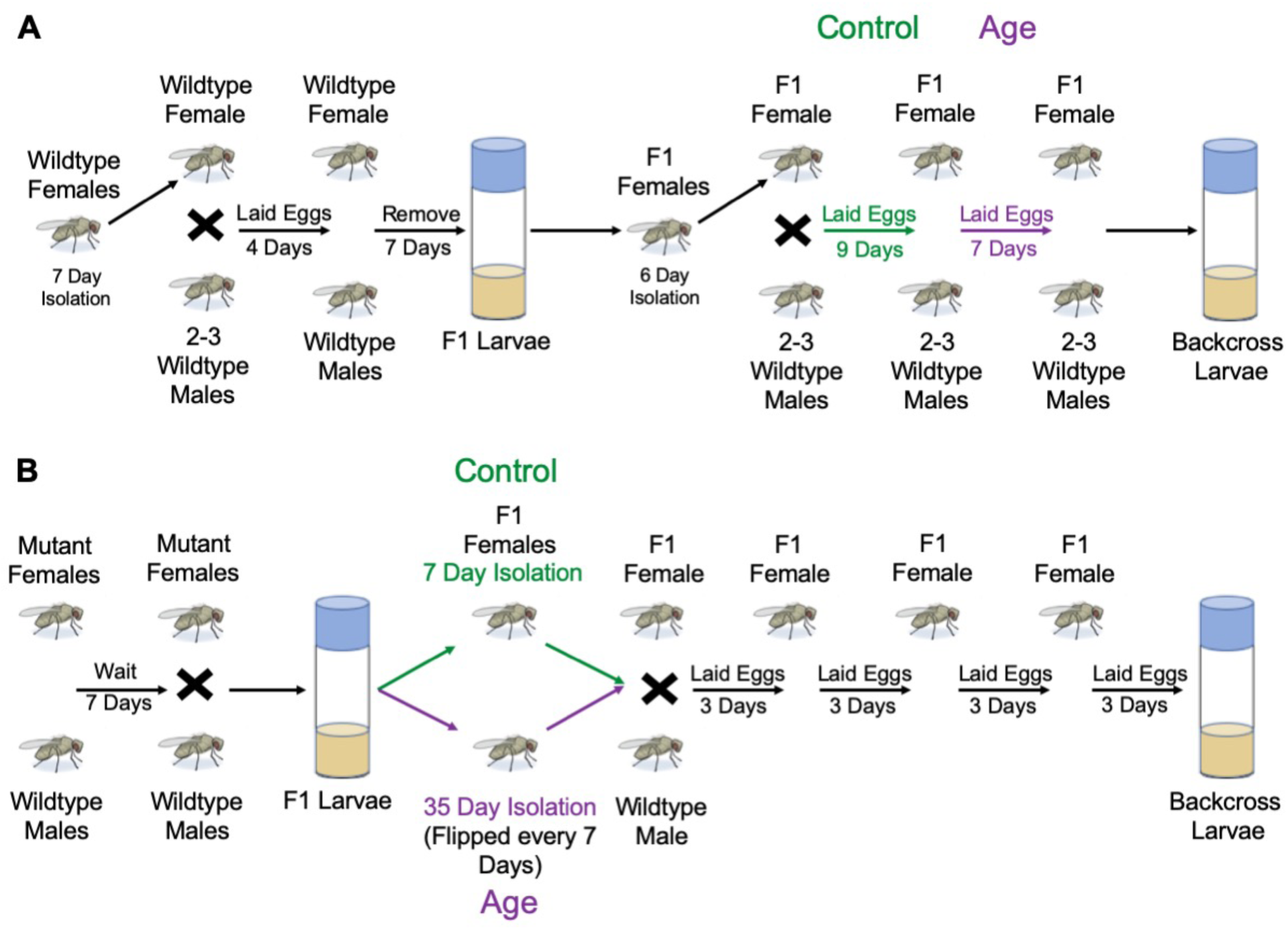
Experimental design. An earlier SNP genotyping study did not find any difference in recombination rate that could be attributed to maternal age in *D. pseudoobscura*. The experimental design (A) of Manzano-Winckler *et al*. (2013), was carefully reviewed in the experimental design for this study (B). While Drosophila lay eggs continuously, artificial clutches can be made by transferring females to a fresh food vial during egg laying. A key difference in this study was that the age difference was distinct and large rather than comparing clutches approximately 1 week apart. This allowed the use of different flies for treatment and control rather than using the same flies for both.

Here, we chose an experimental design that compared smaller clutches of progeny (3 days each) between crosses of females from two distinct maternal ages, with progeny collection continuing for 12 days after mating (Figure 1B). We first conducted two experiments to determine a suitable age for comparison, focusing on how fecundity and survival change with age. Previous work has indicated that the stress level needed to study recombination rate plasticity must be close to the stress that leads to death (Parsons 1988). A pilot experiment on fecundity across a range of ages found no significant difference in females aged 7-35 days of age (Figure 2A). Next, a survivorship analysis revealed females aged 35 days had an average survivorship of 59% (Figure 3), which was selected for the recombination analysis. Then, a large-scale recombination experiment compared recombination rate between six replicate crosses each of females aged to 7 days, which was used as a control age, and the selected 35-day maternal age treatment (Figure 1B). Recombination rate in two intervals along the X chromosome was estimated using three X-linked mutant markers. A significant decline in fecundity due to maternal age was found among the progeny from this experiment (Figure 2B). There was also a significant increase in recombination rate in the maternal age crosses as compared to the control crosses in days 1-3 along one of the two genomic intervals (Figure 4A-B; Figure 5; Table 1). Finally, we investigated crossover interference and did not find any statistical difference due to maternal age (Figure 4C; Table 2), but did notice a reduced mean in the age treatment in days 1-3, suggesting a relaxation in crossover control that explained the recombination results. Although not significant, the last time point also showed an increased mean recombination rate in the control crosses for this genomic interval that corresponds to a decrease in mean crossover interference in control crosses for that time point. These results reflect the complexity of direction of change in recombination rate and interference due to maternal age seen in other taxa.

**Figure 2.**
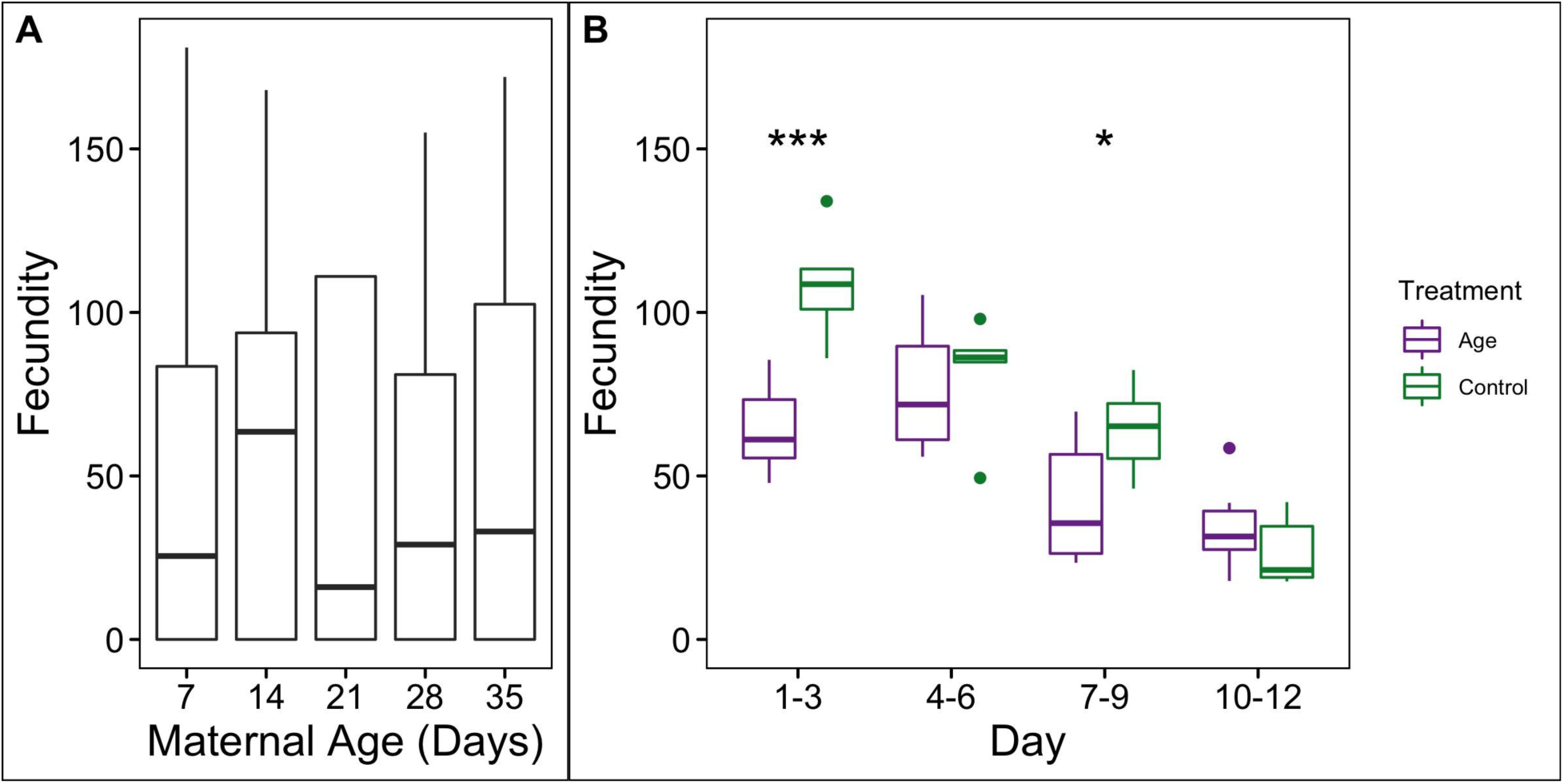
Impact of maternal age on fecundity. A box plot showing the total fecundity across five maternal age treatments and 39 replicate crosses (A). A box plot showing total progeny per backcross per 3-day egg collection period over the 12-day recombination analysis experiment (B). Each time point has a separate box plot for the replicate crosses from the 7-day control (green) and 35-day maternal age treatment (purple). Each F_1_ replicate cross had multiple individual backcrosses which are averaged. Significance between treatment and control for each time point are indicated by asterisks (see Supplementary Table 2).

**Figure 3.**
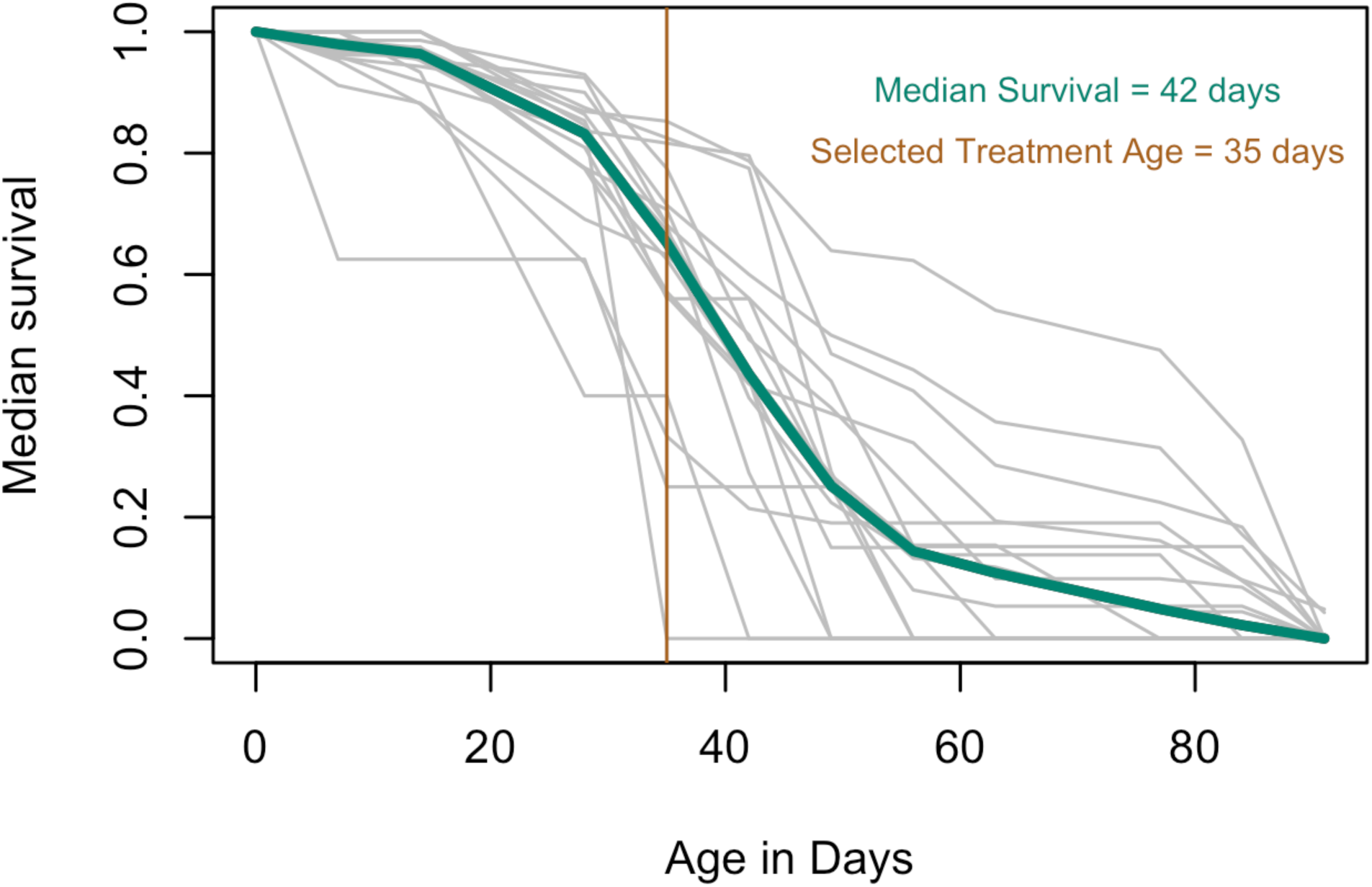
Survivorship declines with age. Percentage of F_1_ females remaining as a function of time. Eighteen replicate crosses were tracked until none were remaining (each shown in grey). Median survivorship for each time point across replicates is shown in dark cyan. The selected treatment for maternal age (dark orange) was the time point immediately preceding the time point where median survival fell below 50%.

**Figure 4.**
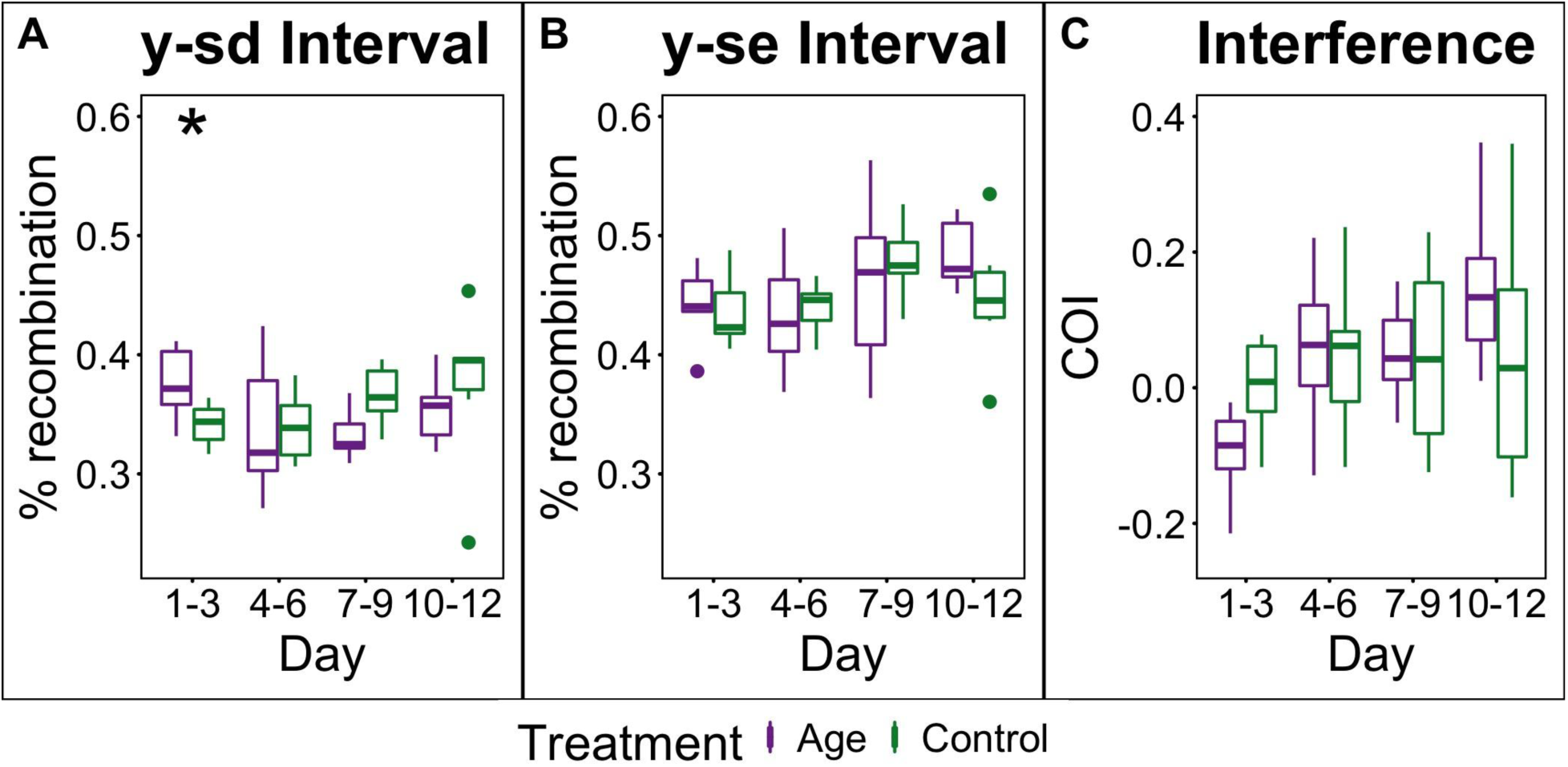
Maternal age alters recombination rate and crossover interference. Recombination rates per time point for 7-day control crosses (green) and crosses with females aged to 35 days (purple) were compared from the *sd-y* region (A) and the *y-se* region (B). Each box plot represents variation among independent F_1_ replicate crosses for each treatment and each time point. Additionally, crossover interference was calculated for each replicate from each treatment and time point (C). Coincidence was defined as the number of double crossovers observed between mutant markers *sd-y-se* versus the number expected based on rates of single crossovers in the intervals *sd-y* and *y-se*. Crossover interference (COI) is coincidence subtracted from 1. Significance between treatment and control for each time point in each plot are based on *post hoc* means contrasts and indicated by asterisks (see Supplementary Tables 3-4).

**Figure 5.**
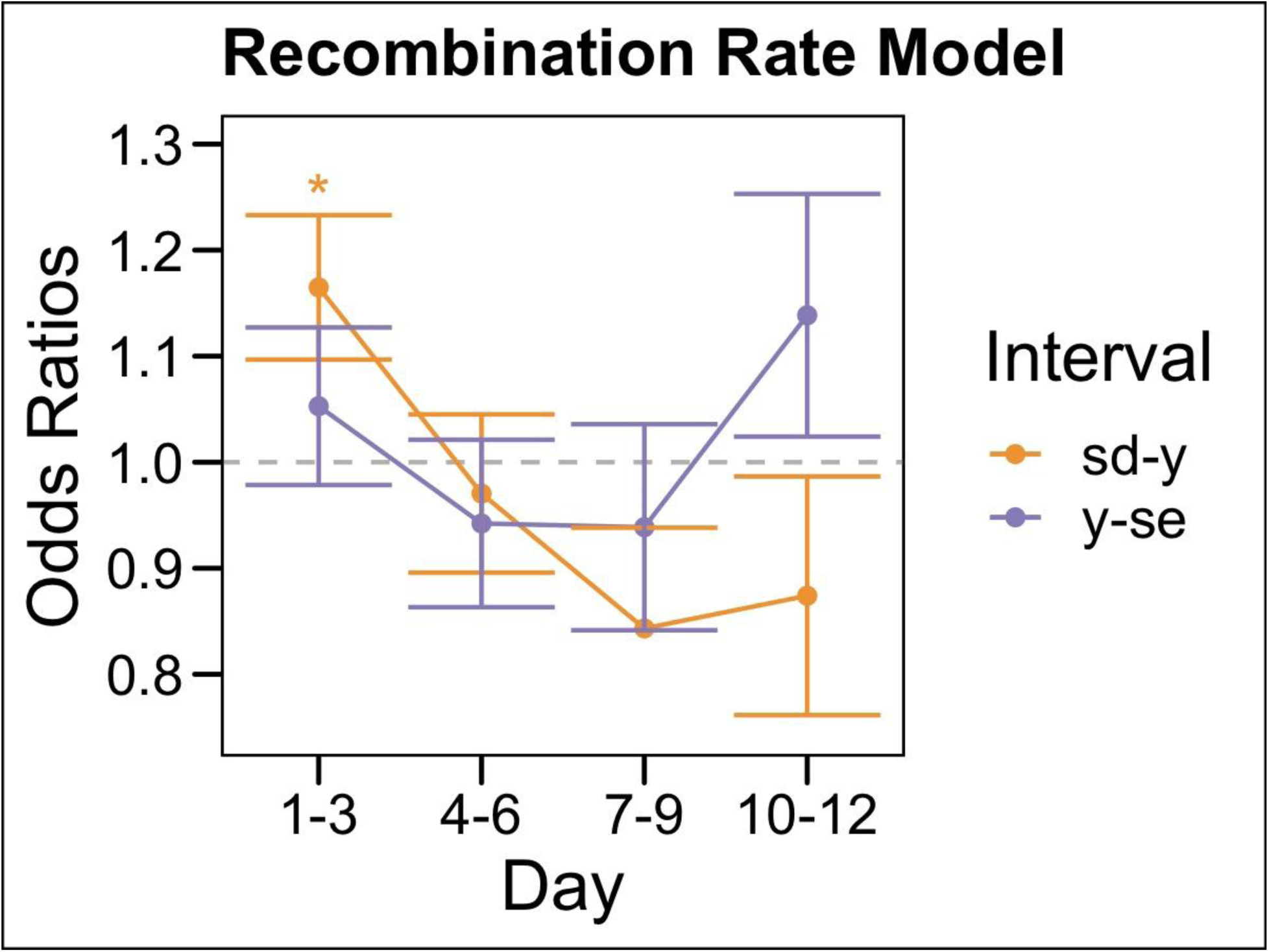
Recombination rate for the *sd-y* interval differs significantly on days 1-3. Recombination rate between control and maternal age were compared using a generalized mixed-effects model in R. Exponentiating the coefficients generated the odds ratio. A positive odds ratio indicates a higher probability of observing a crossover in the maternal age treatment as compared to the control. A *posthoc* test was done to calculate significance for each timepoint between treatment and control. Odds ratio vs. days post-mating are shown for maternal age for the interval between *scalloped* and *yellow* (orange) and for the interval between *yellow* and *sepia* (purple).

**Table 1.**
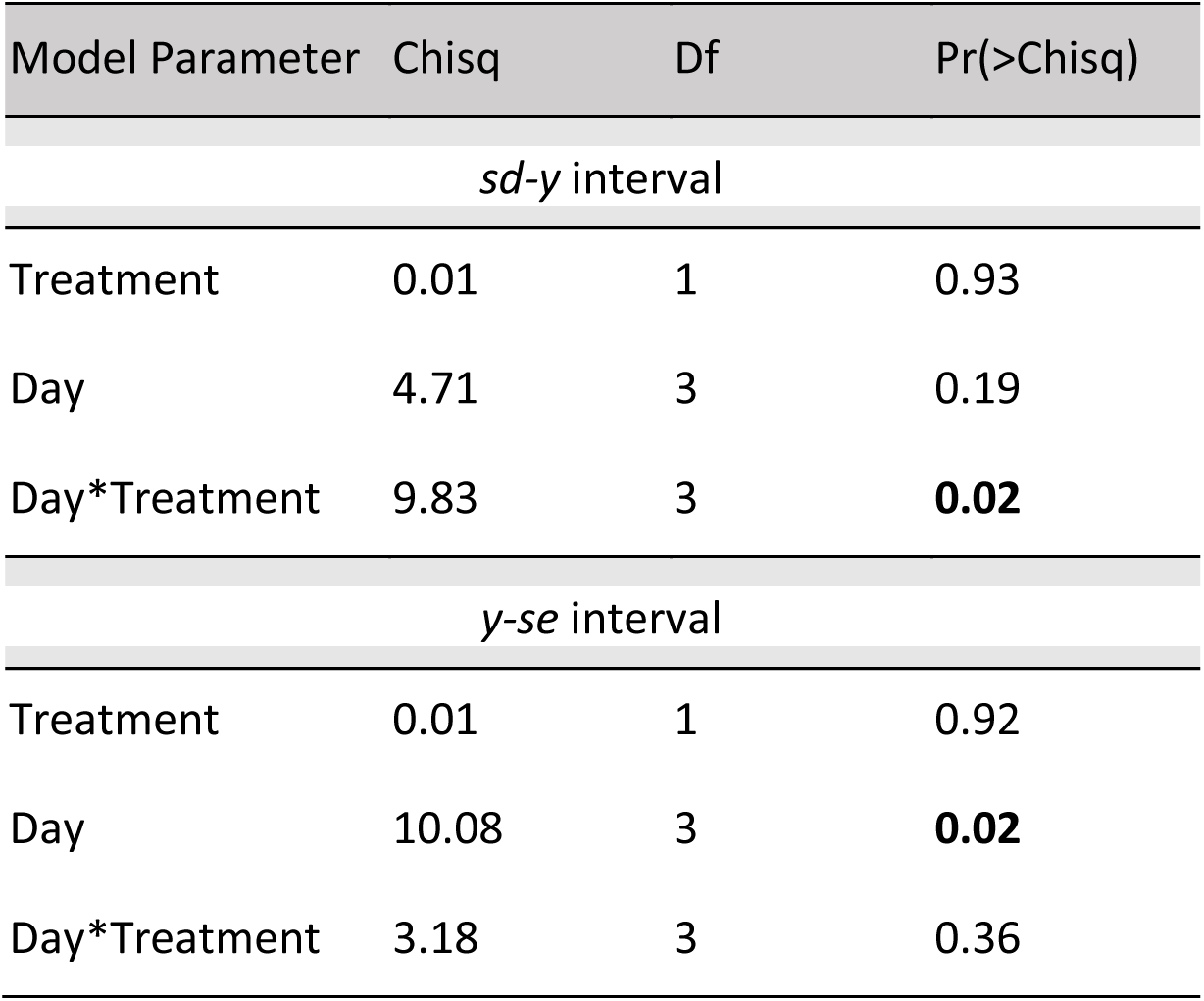
Model Table for Recombination Rate Analysis. A mixed model analysis per interval with replicate IDs as random effects and all other parameters as fixed effects was conducted in R. The model from the GLMer was fit to an Anova table using chi-square with the package ‘car’.

**Table 2.**
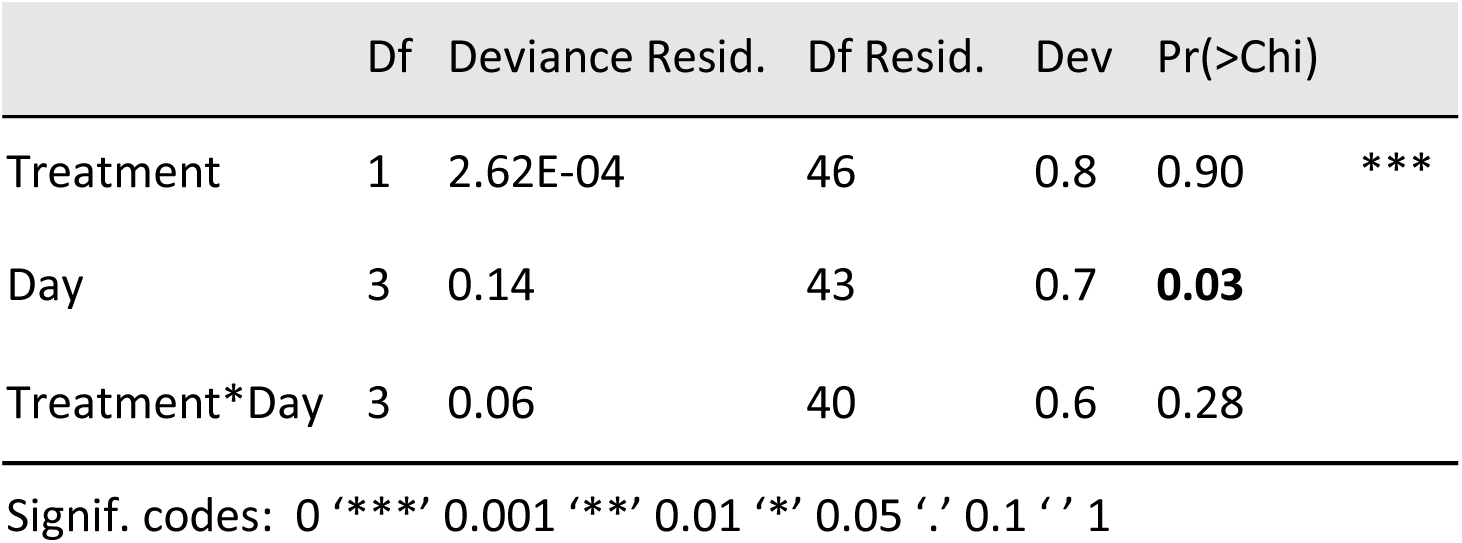
Model Table for Crossover Interference. A linear regression analysis of interference as a response variable and model variables of treatment and day with an interaction term was conducted in R. The resulting glm was fit to an Anova table using chi-square to generate the model table.

## Materials and Methods

### Fecundity Pilot Analysis

Virgin wildtype females from the MV2-25 stock were collected for five weeks and aged to maternal ages ranging from 7-35 days. All crosses were reared in a 21°C incubator on cornmeal-sugar-agar-yeast media. After aging, virgin females were crossed in a 1:1 ratio to wildtype males from the same stock (5-7 days old). After 24 hours, the male was removed to prevent male harassment. As a control measure, the enclosures where virgin females were aged, regardless of treatment, were kept for 14 days after removal of females for use in crosses, to ensure there were no larvae. After 7 days, the female was removed. Once the progeny began to eclose, the number of males and females were recorded each day to calculate fecundity. A total of 1998 progeny were collected from 39 replicate crosses across the maternal ages of 7 (N=10), 14 (N=10), 21 (N=5), 28 (N=7), or 35 (N=7) days. An analysis of variance was conducted in R to test for a difference in fecundity across the five selected age treatments.

**Table 3.** Post hoc table for recombination rate. Means contrast using emmeans package and Kenward-Roger mode were conducted on the models in Table 1 to compare treatment and control for each time point individually.

| contrast | Day | estimate | SE | df | z.ratio | p.value |
| --- | --- | --- | --- | --- | --- | --- |
| *sd-y interval* |  |  |  |  |  |  |
| Age - Control | 1-3 | 0.15 | 0.07 | Inf | 2.24 | **0.025** |
| Age - Control | 4-6 | -0.03 | 0.07 | Inf | -0.40 | 0.688 |
| Age - Control | 7-9 | -0.17 | 0.09 | Inf | -1.80 | 0.073 |
| Age - Control | 10-12 | -0.13 | 0.11 | Inf | -1.20 | 0.232 |
| *y-se interval* |  |  |  |  |  |  |
| Age - Control | 1-3 | 0.05 | 0.07 | Inf | 0.69 | 0.489 |
| Age - Control | 4-6 | -0.06 | 0.08 | Inf | -0.75 | 0.451 |
| Age - Control | 7-9 | -0.06 | 0.10 | Inf | -0.65 | 0.515 |
| Age - Control | 10-12 | 0.13 | 0.11 | Inf | 1.13 | 0.257 |

### Survivorship Analysis

In order to determine longevity of *D. pseudoobscura*, F_1_ females were generated using the same crossing scheme described below for the recombination rate estimates. Eighteen replicate crosses of 10 mutant females with 5 wildtype males were conducted, and the F_1_ female progeny were collected. Progeny were kept in vials with an average of 6.5 females (ranging from 1-13) based on when they were collected. To ensure the females had fresh food supply throughout the experiment, they were transferred to fresh food every 7 days. At each transfer, the number of females remaining in the vial was counted and recorded until no flies were left. For each replicate and time point, the percentage remaining as compared to the initial count was computed. The median across each time point was then computed to identify the time point at which less than 50% females remained. This analysis was used to justify the choice of age selected for the large-scale recombination analysis.

### Recombination Rate Analysis using multiple phenotypic markers

Stocks used, fly husbandry, and cross design are as described in Altindag *et al*. (2020). Briefly, to measure differences in recombination rate, crosses were conducted using a wild type *D. pseudoobscura* stock (stock 14011-0121.265, Dpse\wild-type “TL”, SCI_12.2) and an X-linked recessive mutant *D. pseudoobscura* stock. Recombination rate was measured in two genomic intervals using three phenotypic mutant markers (see Figure 1B in Altindag *et al*. 2020) - *scalloped* (*sd*; 1–43.0), *yellow* (*y*; 1–74.5), and *sepia* (*se*; 1–156.5) (Phadnis 2011). Mutations in the *scalloped* gene alter the wing phenotype, mutations of the *yellow* gene alter the body color, and mutations in the *sepia* gene result in brown eyes.

Crosses were conducted as described in and as shown in Figure 2B in Altindag *et al*. (2020), and correspond closely to the “Experiment 4” design. Briefly, to measure how maternal age impacts recombination rates, homozygous recessive mutant females aged to 7 days were crossed to age-matched wildtype males. The heterozygous F_1_ females were aged to 7 days (control) and 35 days (maternal age treatment) and backcrossed to wild type males to ensure no fitness effects associated with recessive mutant alleles. This cross design also provided a built-in ‘fail safe’ because female progeny could not be homozygous for the recessive mutant markers, and thus any mutant females would be an indicator of contamination (see below). As shown in Figure 1B, the F_1_ females for the maternal age treatment were transferred into new vials every 7 days until they were 35 days old. The collection, crossing, and F_1_ collection of the control flies were timed so they would be backcrossed at the same time as the 35-day old maternal age treatment flies. When the maternal age treatment females were 35 days old and the control females were 7 days old, they were backcrossed to wildtype males.

For both the maternal age treatment and control females, 8-10 backcrosses of females from 6 replicate F_1_ crosses were conducted. The wildtype males were age matched to the control (7 days old) and isolated 24 hours before genetic crosses. To backcross, a single wildtype male and F_1_ female were placed in a fresh food vial. To promote mating, a cotton ball was placed inside to restrict available space and the vial was placed under a 100 Watt CFL light for an hour. Afterwards, they were returned to the incubator with a 12 hour light-dark cycle. After 24 hours, the cotton was removed and the male was discarded to restrict crosses to a single mating. The female continued to be transferred to a fresh food vial every 3 days for 12 days (Figure 1B). When the progeny begin to eclose, they were transferred into a new vial and allowed to age 5 days prior to phenotyping to allow the *sepia* eye color mutation to become more prominent. This delayed phenotyping ensured accuracy in scoring progeny. Because the markers were X-linked recessive and backcrosses were to wild type males, only male progeny were scored and if any female progeny were found to be mutant, the entire vial was discarded and the data removed (see above). During progeny collection, each of the three traits were recorded as either mutant or wild type independently in a single-blind manner. Each pair of markers allowed for recombination rate estimates to be calculated between the *scalloped* and *yellow* markers (hereafter *sd-y* interval) and between the *yellow* and *sepia* markers (hereafter *y-se* interval). Phenotyping ended two weeks after eclosion started to prevent the next generation from being included in the data. Every effort was made to score all progeny. Data were entered in triplicate and compared until 100% concordant.

### Crossover interference

The occurrence of a crossover in one genetic region is usually associated with a decreased probability of a concomitant crossover in an adjacent region. This phenomenon is called positive crossover interference (COI). In this study, the region *sd-y-se* allowed for the estimation of expected double crossover frequencies. COI was calculated by dividing the observed frequency of double recombinant progeny to the expected frequency of double recombinants. Across the replicates for both control and the maternal age treatment, the expected number of double crossover events for the *sd-y-se* region was calculated by the product of individual single crossover frequencies in each interval (*sd-y* and *y-se*) and for each time point across the 12 day experiment.

### Statistical Analysis of Recombination Data

Statistical analysis was performed using R v4.0.1 (R Core Team 2020). R was used to conduct mixed-model ANOVA in order to identify the difference in recombination rates between experimental and control conditions following the methods outlined in (Altindag *et al*. 2020). The R code used for all statistical analysis and to produce all data figures and model tables is available on github (see below). While 8-10 backcrosses were conducted per replicate, this was done mainly to reduce male harassment by limiting females to a single mating event. Therefore, the sample sizes from these crosses were averaged across backcrosses for each independent F_1_ replicate cross for the purposes of analysis. For crossover events, the results were summed across backcrosses for each independent F_1_ replicate cross.

First, to investigate the impact of maternal age on fecundity throughout the experiment, the number of progeny over the 12 day experiment was treated as the response variable. A quasipoisson regression was conducted in R (Supplementary Tables 1 and 2). Next, crossover counts were analyzed in a logistic regression analysis for each genomic interval as done in Altindag *et al*. (2020). Third, the COI results were analyzed using a general linear model. For each of the three analyses, model parameters were treatment, time point, and the interaction, and a *posthoc* means contrast was subsequently conducted to examine the differences between treatment and control for each time point.

## Data Availability

Data files and scripts to complete the analysis are available on github at https://github.com/StevisonLab/Maternal-Age-Project. A snapshot of the code at the time of publication is available at DOI: 10.5281/zenodo.3951055. This repository includes the pilot fecundity data, raw survivorship data, raw mutant phenotype records for males, and female count data from the recombination analysis are included as csv files along with the code. A separate csv file with treatment information includes dates and other metadata that would be needed to validate the analysis and conclusions herein. Additionally, a processed data file that includes sums of males and females, as well as crossover counts across each interval per time point, per replicate cross is also included.

## Results

### Fecundity does not vary across selected age treatments

Progeny from 39 replicates across the maternal ages ranging from 7-35 days were collected to determine if maternal age affected fecundity (Figure 2A). These pilot data showed an average fecundity per replicate of 51.23 progeny. The average fecundity was highest for the 14-day females (60 progeny per replicate), and lowest for the 21 day females (47.6 progeny per replicate). An analysis of variance revealed no significant difference in fecundity across the selected maternal age treatments (p=0.988). Still, the 35-day and 28-day treatment had a higher rate of replicates producing no offspring (43%) compared to the other treatments (40%).

### Median survivorship is 42 days of age

To determine the longevity of this species, 808 progeny across 18 replicates were followed until none remained (91 days). Each replicate cross had an average of 46.28 females ranging from 8 to 75 (housed in smaller groups, see Methods). The percentage of survival decreased slowly between day 0 and day 30, rapidly between day 30 and day 57, and slowly between day 57 and day 91 (Figure 3). The inflection point of the survivorship curve where the median survivorship was below 50% occurred at 42 days. At 42 days, 61% of replicates had more than 50% loss as compared to original counts. The time point immediately preceding this was 35 days. At 35 days, only 22% of replicates had greater than 50% loss. Therefore, 35 days was selected as the treatment for the mutant screen experiment to investigate how maternal age impacts recombination rate.

### Maternal age alters recombination rate and fecundity

To measure recombination rates, phenotypes at three mutant X-linked markers were recorded for 10,727 male flies (n_age_=4,508; n_control_=6,219). The mean total recombination rate for the control samples was 80.6%. The recombination rate between *yellow* and *sepia* for the control crosses was 45.1%, which was different from the expected map distance (81cM), but similar to the rate found in a previous experiment conducted with a similar experimental design (Altindag *et al*. 2020; 46.0%). Note that markers greater than 50cM apart will not recover recombination frequencies above 50%. Between *yellow* and *scalloped*, the mean recombination rate for the control crosses was 35.5%. This is different, but close to the expected map distance (32.5 cM), and similar to the recombination rate found in the same interval in a previous experiment (Altindag *et al*. 2020; 38.5%).

During progeny collection, fecundity was tracked to compare the control and maternal age treatment (Figure 2B). The total number of progeny collected was 23,559 (n_age_=10,050; n_control_=13,509). Unlike the pilot results on fecundity, there was a significant difference in fecundity between females aged 7 days used for the control crosses (mean=70.36) and females aged to 35 days (mean=54.29). Similar to previous work (Altindag *et al*. 2020), fecundity declined steadily throughout progeny collection, consistent with a single mating event. The fecundity analysis showed a significant effect of treatment on fecundity (p=9.34E-4). A *posthoc* mean contrast found that fecundity was significantly different between treatments for the 1-3-day time point (p=1.46E-4) and the 7-9-day time point (p=0.013).

The statistical comparison of recombination rate between the maternal age treatment and control did not show a significant effect of treatment on recombination rate for either interval (Table 1). However, for the *sd-y* interval, the interaction between time point and treatment was significant (p=0.02). Additionally, there was a significant effect of time point for the *y-se* interval (p=0.02). The odds ratio (OR) between maternal age treatment and control were exponentiated from the model coefficients (Figure 5). A *posthoc* mean contrast analysis revealed a significant difference in recombination rate (p=0.025; OR=1.16) in the first 72 hour time point for the *sd-y* interval (starred in Figure 5; Supplementary Table 3). Though results were not significant for the *y-se* interval, it also shows a higher mean recombination rate in the first time point for maternal age (Figure 4A-B). Additionally, the *y-sd* interval shows higher recombination rate in control crosses for later time points, though these results are not statistically significant (see Discussion).

### Reduced crossover interference due to maternal age

The calculated COI across control and maternal age replicates did not show any difference due to treatment (Table 2). However, time point was significant in the overall model (p=0.03), indicating the COI varied across the experiment. Though not significant, it is worth pointing out that the COI is lower in the maternal age replicate crosses as compared to the control crosses for the first time point (Figure 4C), which can explain the significant increase in recombination rate for the time point (see Discussion). Additionally, the last time point has higher COI in the maternal age treatment than the control, closely matching the recombination rate data.

## Discussion

Results indicated a significant increase in recombination rate in days 1-3 post-mating (3.39%; starred in Figure 4A) due to maternal age in *D. pseudoobscura*. Interestingly, these results correspond to a significantly lower fecundity (Figure 2B), and lower crossover interference (Figure 4C) in the maternal age treatment at the same time point. These results contrast previous work which failed to find an effect of maternal age on recombination rate in this species (Manzano-Winkler *et al*. 2013). However, this difference was likely due to differences in experimental design (see Introduction; Figure 1A), and increased sample size. Additionally, it is worth noting that the previous study compared recombination rate between three regions on chromosome 2 using molecular markers, in contrast to this study which compared recombination between two regions of the X chromosome using phenotypic mutants. Therefore, it is also possible that the genomic regions vary in their response to maternal age (Grell 1978).

The results here differed from recent experiments on recombination rate plasticity due to thermal stress (Altindag *et al*. 2020). In a companion study, differences in recombination rate due to temperature peaked at 9 days post-mating, and had a much larger effect size (10.97% increase at 26°C as compared to 21°C), even with a smaller sample size. This difference in timing and magnitude of effect suggests a distinctly different mechanism for maternal age in recombination rate plasticity than temperature. It is possible that the temperature and maternal age impact different time points in early meiosis, which leads to the difference in timing between experiments. Alternatively, heat stress could lead to delays in timing of early meiotic events (Joyce and McKim 2010; Draeger and Moore 2017), which would lead to an apparent shift in when the recombination rate difference is observed in the resulting progeny. Perturbation experiments would be able to distinguish these two possibilities (Grell 1973; Saini *et al*. 2017). Reducing the transfer frequencies from 72 hours to 24 hours would allow for the timing of maternal age plasticity to be further narrowed prior to perturbation experiments (Altindag *et al*. 2020).

**Table 4.** Post hoc table for crossover interference. Means contrast using emmeans package and kenward-roger mode were conducted on the model in Table 2 to compare treatment and control for each time point individually.

| contrast | Day | estimate | SE | df | z.ratio | p.value |
| --- | --- | --- | --- | --- | --- | --- |
| Control - Age | 1-3 | 0.10 | 0.07 | Inf | 1.36 | 0.17 |
| Control - Age | 4-6 | -0.01 | 0.07 | Inf | -0.13 | 0.89 |
| Control - Age | 7-9 | -0.01 | 0.07 | Inf | -0.08 | 0.94 |
| Control - Age | 10-12 | -0.10 | 0.07 | Inf | -1.41 | 0.16 |

### Fecundity and survivorship results support reduced fitness due to maternal age

Reduced natural selection later in life has been implicated for age-related fitness declines, such as fecundity and longevity. The fecundity and survivorship results here were consistent with decreased fitness with maternal age. Fertility of *Drosophila* depends on successful oogenesis, ability to store sperm, fertilization success, and the age of the parents. In *D. melanogaster*, oogenesis is affected by maternal age and the number of oocytes produced decreases as the age of the mother increases (Brenman 2017). Additionally, a 50% decline in fecundity is observed between mothers aged 22-28 days old and mothers aged 4-7 days old in *D. melanogaster* (Miller *et al*. 2014). Although the pilot experiment here did not detect a significant difference due to age (Figure 2A), the ages selected may not have ranged enough to detect any difference. In the survivorship analysis, females lived up to 91 days, and it was not until 42 days of age that longevity reduced below 50%. Fecundity in the recombination analysis showed the expected significant decrease (22.84%) between 35-day old and 7-day old females (Figure 2B). Additionally, the survivorship results here (Figure 3) were consistent with a type 2 survivorship curve, which represents increased variation in age at death among individuals, as compared to type 1 and type 3 curves (Demetrius 1978).

### Direction of response in recombination and COI shifts over time

In work done by Bridges (1927) and Redfield (1966) in *D. melanogaster* an M-shape or “rise-fall-rise” was pattern observed for recombination rate. This seems to be somewhat driven by changes in interference, which havs an inverse pattern, or W-shape response (Bridges 1927). Here, there was a significant increase in recombination rate in days 1-3 due to maternal age, and for days 7-12, age had a lower mean recombination rate (n.s.), shared across both intervals (Figure 4A-B). Similarly, the maternal age treatment had lower COI in days 1-3, but a higher mean COI in days 10-12 (Figure 4C). A lower COI suggests relaxation in the regulatory control of crossovers, which has also been shown in humans and plants to decrease with age (Toyota *et al*. 2011; Campbell *et al*. 2015). However, the effect of interference on crossover formation varies across the genome and differences in both sex and age result in variation of recombination events depending on the spatial distribution of alleles (Grell 1978; Toyota *et al*. 2011).

Additionally, there was more variation in recombination rate across time points in the maternal age replicates than in the control replicates. In contrast, for COI, there was more variation across time points in the control replicates than in the maternal age replicates. This increased variance may contribute to the overall small effect size observed. Indeed, a very large sample size was necessary to capture the differences in recombination rate due to maternal age observed here.

## Acknowledgements

This work was supported by research start-up funds to LSS from the Department of Biological Sciences at Auburn University. We thank the Phadnis Lab for supplying the mutant stock used for this work. We thank Todd Steury for consultation on our statistical analysis. We thank members of the Stevison Lab for extensive help with recording mutant marker phenotypes, with particular thanks to Anna Tourne, Kaitlyn Walter, and several other undergraduate researchers.

## 100 word article summary

Fertility and longevity, which affect an organism’s fitness, decline with age. Aging also alters recombination rates, which can increase meiotic errors. Here, the effect of maternal age on recombination rate was investigated in *Drosophilapseudoobscura* via a large-scale genetic cross. Recombination frequency was 3.39% higher within 72 hours of mating in females aged to 35 days as compared to 7-day old females, with corresponding decreases in fecundity and crossover interference. These findings contrast studies on temperature induced recombination rate plasticity, which differ in recombination later, suggesting different mechanisms mediate the effect of temperature and maternal age on recombination rate.

## Data Availability

Github Repository: https://github.com/StevisonLab/Maternal-Age-Project

DOI: 10.5281/zenodo.3951055

## Supplementary Tables

**Table S1.**
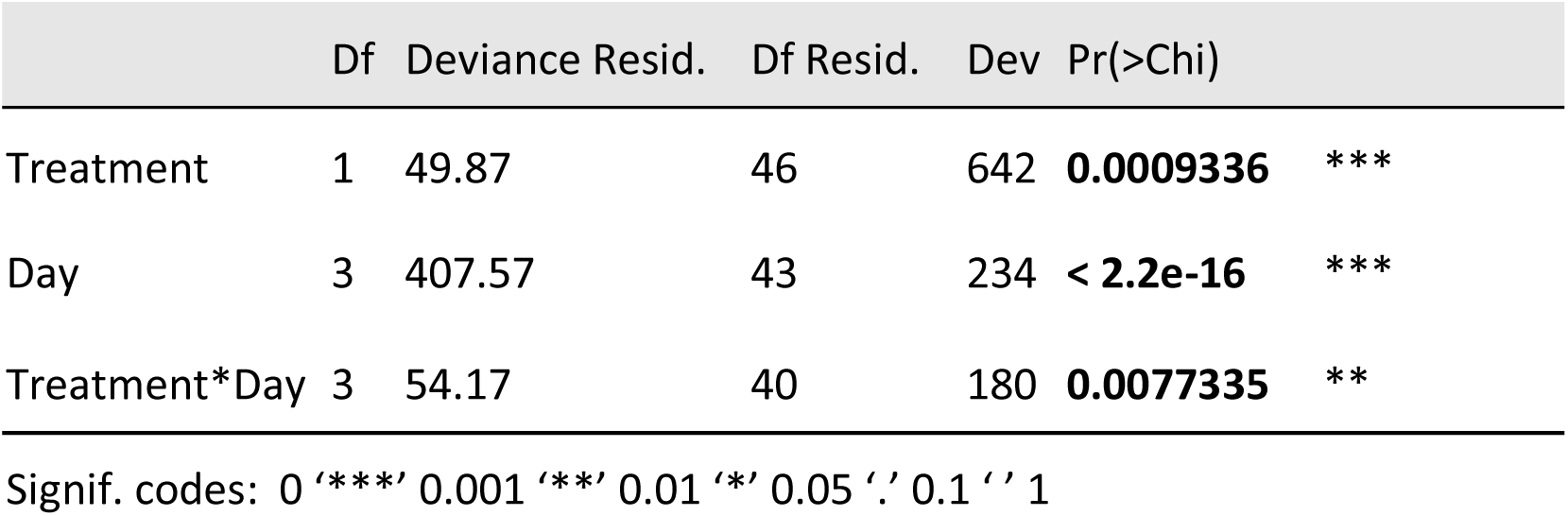
Model Table for Fecundity analysis in large-scale mutant screen. Quasipoisson regression analysis of fecundity as a response variable and model variables of treatment and day with an interaction term was conducted in R. The resulting glm was fit to an Anova table using chi-square to generate the model table.

**Table S2.** Post hoc table for fecundity in large-scale mutant screen. Means contrast using emmeans package and Kenward-Roger mode were conducted on the model in Supplementary Table 1 to compare treatment and control for each time point individually.

| contrast | Day | estimate | SE | df | z.ratio | p.value |
| --- | --- | --- | --- | --- | --- | --- |
| Control - Age | 1-3 | 0.52 | 0.14 | Inf | 3.80 | **1.46E-04** |
| Control - Age | 4-6 | 0.07 | 0.14 | Inf | 0.54 | 0.59 |
| Control - Age | 7-9 | 0.43 | 0.17 | Inf | 2.48 | **1.32E-02** |
| Control - Age | 10-12 | -0.26 | 0.22 | Inf | -1.17 | 0.24 |

## References

Agrawal, A. F., L. Hadany, and S. P. Otto, 2005 The Evolution of Plastic Recombination. Genetics 171: 803–812.

Altindag, U. H., C. Shoben, and L. S. Stevison, 2020 Refining the timing of recombination rate plasticity in response to temperature in *Drosophila pseudoobscura*. bioRxiv 2020.04.10.036129.

Brenman, D., 2017 The Effect Of Age On Social Behaviour In Drosophila melanogaster And The Progeny of Aged Parents: The University of Western Ontario, 158 p.

Bridges, C. B., 1915 A linkage variation in Drosophila. J. Exp. Zool. 19: 1–21.

Bridges, C. B., 1927 The Relation Of The Age Of The Female To Crossing Over In The Third Chromosome Of *Drosophila melanogaster*. J. Gen. Physiol. 8: 689–700.

Campbell, C. L., N. A. Furlotte, N. Eriksson, D. Hinds, and A. Auton, 2015 Escape from crossover interference increases with maternal age. Nat. Commun. 6: 6260.

Charlesworth, B., 2000 Fisher, Medawar, Hamilton and the Evolution of Aging. Genetics 156: 927–931.

Draeger, T., and G. Moore, 2017 Short periods of high temperature during meiosis prevent normal meiotic progression and reduce grain number in hexaploid wheat (*Triticum aestivum L.*). TAG Theor. Appl. Genet. Theor. Angew. Genet. 130: 1785–1800.

Felsenstein, J., 1974 Evolutionary Advantage of Recombination. Genetics 78: 737–756.

Flatt, T., and P. S. Schmidt, 2009 Integrating evolutionary and molecular genetics of aging. Biochim. Biophys. Acta 1790: 951–962.

Grell, R. F., 1978 A Comparison of Heat and Interchromosomal Effects on Recombination and Interference in *Drosophila melanogaster*. Genetics 89: 65–77.

Grell, R. F., 1973 Recombination and DNA replication in the *Drosophila melanogaster* oocyte.22.

Hales, K. G., C. A. Korey, A. M. Larracuente, and D. M. Roberts, 2015 Genetics on the Fly: A Primer on the *Drosophila* Model System. Genetics 201: 815–842.

Hassold, T., and P. Hunt, 2001 To err (meiotically) is human: the genesis of human aneuploidy. Nat. Rev. Genet. 2: 280–91.

Henderson, S. A., and R. G. Edwards, 1968 Chiasma Frequency and Maternal Age in Mammals. Nature 218: 22–28.

Joyce, E. F., and K. S. McKim, 2010 Chromosome Axis Defects Induce a Checkpoint-Mediated Delay and Interchromosomal Effect on Crossing Over during *Drosophila* Meiosis (G. P. Copenhaver, Ed.). PLoS Genet. 6: e1001059.

Kong, A., J. Barnard, D. F. Gudbjartsson, G. Thorleifsson, G. Jonsdottir et al., 2004 Recombination rate and reproductive success in humans. Nat. Genet. 36: 1203–1206.

Manzano-Winkler, B., S. E. McGaugh, and M. A. F. Noor, 2013 How Hot Are Drosophila Hotspots? Examining Recombination Rate Variation and Associations with Nucleotide Diversity, Divergence, and Maternal Age in *Drosophila pseudoobscura*. PLOS ONE 8: e71582.

Miller, P. B., O. T. Obrik-Uloho, M. H. Phan, C. L. Medrano, J. S. Renier et al., 2014 The song of the old mother: Reproductive senescence in female *Drosophila*. Fly (Austin) 8: 127–139.

Parsons, P. A., 1988 Evolutionary rates: effects of stress upon recombination. Biol. J. Linn. Soc. 35: 49–68.

Parsons, P. A., 1962 Maternal age and developmental variability. J. Exp. Biol. 39: 251–260.

Parsons, P. A., 1964 Parental Age and the Offspring. Q. Rev. Biol. 39: 258–275.

R Core Team, 2020 R: A language and environment for statistical computing. R Foundation for Statistical Computing, Vienna, Austria.

Redfield, H., 1966 Delayed Mating and the Relationship of Recombination to Maternal Age in *Drosophila melanogaster*. Genetics 53: 593–607.

Rose, A. M., and D. L. Baillie, 1979 The Effect of Temperature and Parental Age on Recombination and Nondisjunction in *Caenorhabditis elegans*. Genetics 92: 409–418.

Saini, R., A. K. Singh, S. Dhanapal, T. H. Saeed, G. J. Hyde et al., 2017 Brief temperature stress during reproductive stages alters meiotic recombination and somatic mutation rates in the progeny of *Arabidopsis*. BMC Plant Biol. 17: 103.

Smith, J. M., and J. Haigh, 1974 Hitch-Hiking Effect of a Favorable Gene. Genet. Res. 23: 23–35.

Stevison, L. S., S. Sefick, C. Rushton, and R. M. Graze, 2017 Recombination rate plasticity: revealing mechanisms by design. Philos. Trans. R. Soc. B Biol. Sci. 372: 20160459.

Throckmorton, L. H., 1975 The phylogeny, ecology, and geography of *Drosophila*, pp. 421–469 in Handbook of Genetics, edited by R. C. King, Plenum, New York.

Toyota, M., K. Matsuda, T. Kakutani, M. T. Morita, and M. Tasaka, 2011 Developmental changes in crossover frequency in Arabidopsis. Plant J. 65: 589–599.

